# Biantennary N-glycans As Receptors for MARTX Toxins in *Vibrio* Pathogenesis

**DOI:** 10.1101/2024.09.12.611726

**Authors:** Jiexi Chen, Felix Goerdeler, Thapakorn Jaroentomeechai, Francisco X. S. Hernandez, Xiaozhong Wang, Henrik Clausen, Yoshiki Narimatsu, Karla J. F. Satchell

## Abstract

Multifunctional Autoprocessing Repeats-in-Toxin (MARTX) toxins are a diverse effector delivery platform of many Gram-negative bacteria that infect mammals, insects, and aquatic animal hosts. The mechanisms by which these toxins recognize host cell receptors for translocation of toxic effectors into the cell have remained elusive. Here, we map the first surface receptor-binding domain of a MARTX toxin from the highly lethal foodborne pathogen *Vibrio vulnificus*. This domain corresponds to a 273-amino acid sequence with predicted symmetrical immunoglobulin-like folds. We demonstrate that this domain binds internal *N*-acetylglucosamine on complex biantennary N-glycans with select preference for L1CAM and other N-glycoproteins with multiple N-glycans on host cell surfaces. This receptor binding domain is essential for *V. vulnificus* pathogenesis during intestinal infection. The identification of a highly conserved motif universally present as part of all N-glycans correlates with the *V. vulnificus* MARTX toxin boasting broad specificity and targeting nearly all cell types.

## INTRODUCTION

Bacterial toxins rank among the most potent modulators of cellular processes in biology (*1*, *2*). In particular, proteins secreted by bacteria, known as exotoxins, frequently serve as primary virulence factors during infections. They can function at the bacterial colonization site or migrate to distant organs, eliciting a spectrum of pathophysiological responses that result in the predominant symptoms observed in human infection (*1–4*). Studying how these potent disease-causing toxins interact with host cells provides profound insights for developing effective strategies to combat infections and deepen our fundamental understanding of mammalian cellular pathways in host-pathogen interaction. Multifunctional Autoprocessing Repeats-in Toxin (MARTX) toxins are a family of bacterial exotoxins crucial for the pathogenesis of many Gram-negative bacteria (*5–9*). Studies on MARTX toxins from *Vibrio cholerae* (*10–12*), *Vibrio vulnificus* (*13–15*), *Aeromonas hydrophila* (*16*) and others (*17*) have consistently demonstrated their critical roles in facilitating successful colonization and enhancing virulence. These toxins are also well-known for their broad host tropism, enabling them to target a wide range of organisms, from insects and aquatic species to mammals. Their significance is also underscored by their ability to assemble from modular domains with different cytotoxic mechanisms into a single polypeptide array, making them analogous to a ’clustered bomb’ in their effectiveness against host cells (*12*).

MARTX toxins are a unique subfamily of the larger pore-forming Repeats-in-Toxin (RTX) superfamily (*9*). A defining feature of the subfamily is nonclassical C-terminal RTX repeats (“C” repeats), starting with the first nine amino acids (aa) consistent with the RTX nonapeptide repeat motif (GGxG(N/D)Dxx), followed by nine additional amino acids that complete the MARTX C-terminal repeats. To further distinguish from other RTX toxins, MARTX toxins also exhibit conserved N-terminal repeats that can be categorized into “A” or “B” repeats, each harboring an individual amino acid repeat sequence while sharing a central glycine-rich repeat motif of G-7x-G-4x-N (*17*). The region between the “B” and “C” repeats includes a linker region, an effector locus that houses an array of bioactive effectors, and a cysteine protease domain (CPD) (Fig. 1A). Comparing MARTX toxins across different species, the repeat regions, the linker region, and the CPD are conserved in length and aa sequence. In contrast, the effector locus carries a variable number of effectors selected from a repertoire of modular domains shared by the MARTX toxin family members (*17*).

**Fig. 1.**
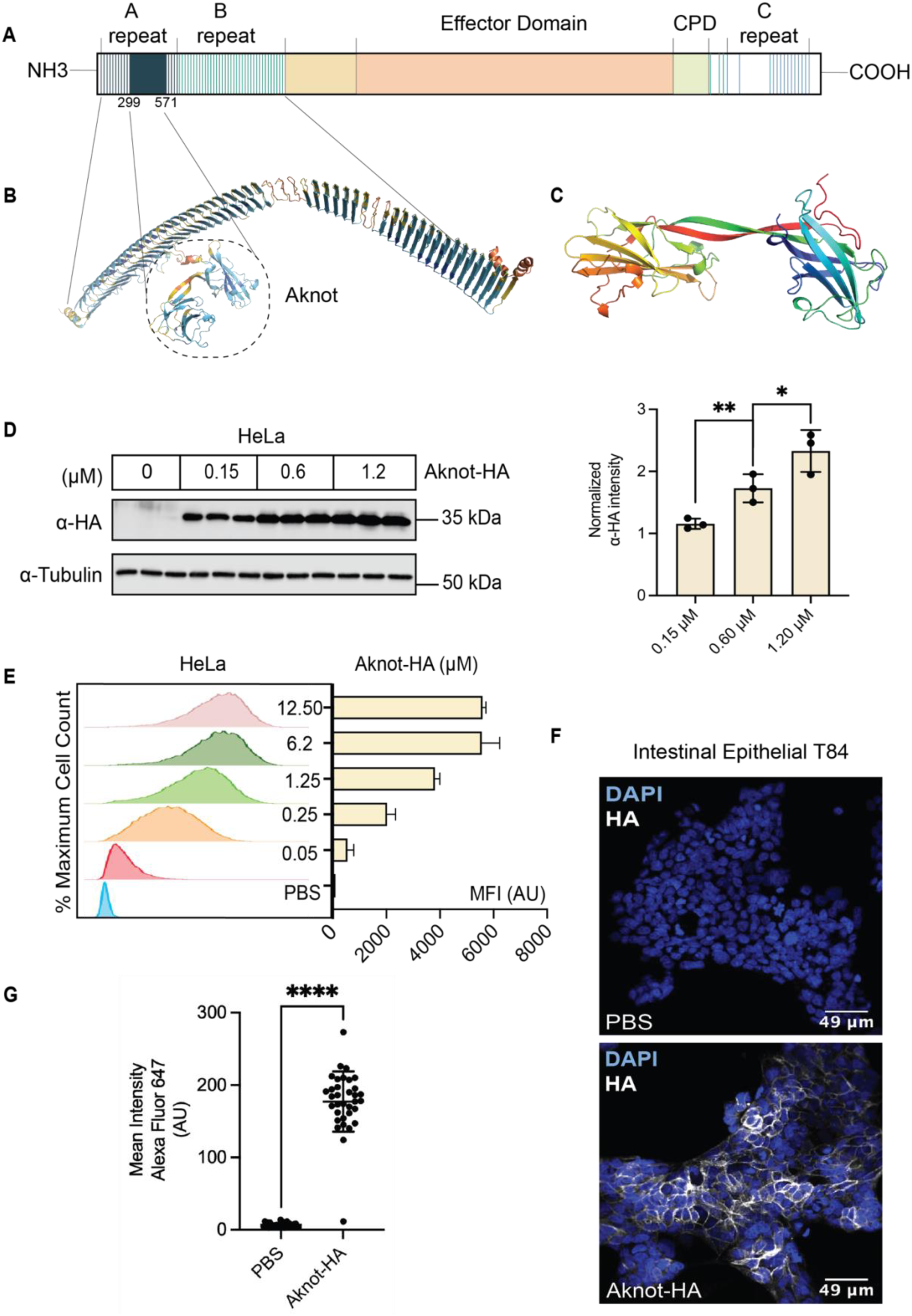
MARTX_Vv_ contains a receptor-binding domain within its N-terminal repeat regions. (A) Schematic representation of the structure organization of *V. vulnificus* MARTX toxin. NH3, protein N-terminus, NH3; CPD, cysteine protease domain, CPD; C-terminus, COOH. Each vertical line represents an individual repeat unit. (B) AlphaFold2 prediction of “A” and “B” repeat regions of MARTX toxin colored by per-residue predicted Local Distance Difference Test (pLDDT) scores. The region circled in dashed lines between the 10^th^ and 11^th^ of “A” repeats, is annotated as “Aknot”. (C) AlphaFold2 prediction of Aknot colored in rainbow gradient from N terminus (red) to C terminus (blue). (D) Representative western blot of Aknot-HA binding to HeLa cells at the indicated concentrations. Three biological replicates are loaded in parallel for each concentration. The quantification of Aknot-HA binding are shown in the bar graph. The signal intensity ratios of Anti-HA/Anti-Tubulin of each concentration are presented as means ± SD from *n=*3 individual experiments. (\**p*<0.05, \*\**p*<0.01, Student’s *t* tests) (E) Representative flow cytometry plots of Aknot-HA binding to HeLa cells and quantification of mean MFI of Alexa Fluor 647 conjugated anti-HA antibody for *n*=3 biological replicates at the indicated concentrations. AU: arbitrary unit. (F) Representative immunofluorescence imaging of 500 nM Aknot-HA binding to intestinal T84 cells. Non-permeabilized cells were stained with Anti-HA for Aknot-HA and DAPI for nuclear staining. (G) Quantification of Aknot-HA binding in (F) from *n*=3 individual experiments analyzed by CellProfiler. (\*\*\*\**p*<0.0001, Student’s *t* tests)

The halophilic Gram-negative bacterium *V. vulnificus* is a marine pathogen that depends on the MARTX toxin for disease pathogenesis. Human *V. vulnificus* infections include seafood-associated gastroenteritis that can progress to sepsis as well as wound infections following contact with seawater (*18*). Although only 100-200 cases are reported annually in the United States, *V. vulnificus* infections have a mortality rate up to 50% and account for 95% of all seafood-related deaths (*19*). The mortality rate can escalate to 100% within 72 hours post-infection in the absence of timely antibiotic treatment (*20*). Notably, an eightfold increase in the incidence of *V. vulnificus* was observed from 1988 to 2018 in the Eastern United States, a trend that strongly correlates with the rise in coastal sea surface temperatures attributable to climate change (*21*). Of particular concern is that *V. vulnificus* infections have emerged in higher latitude areas traditionally considered non-endemic (*22–24*). The MARTX toxins secreted by *V. vulnificus* (MARTX_Vv_) play an indispensable role in the pathogenesis. Mice infected intragastrically with *V. vulnificus* lacking the MARTX encoding gene *rtxA1* exhibit a 180-2600-fold reduction in virulence (*14*). Notably, these mutants fail to spread from the intestine to the bloodstream, linking the MARTX toxin with the progression of the disease to fatal sepsis. Similarly, in wound infection models, the *rtxA1* mutants show a 72-fold reduction in virulence compared to wild-type strains and fail to disseminate from the initial site of infection (*25*, *26*).

Given the significant role of MARTX_Vv_ in the pathogenesis of infections, understanding its mechanism of action is crucial. According to the current conceptual model, MARTX toxins are secreted out of the bacterium by a Type I Secretion System (T1SS) (*27*). The exoprotein then forms a self-assembled pore on the host membrane to enable the translocation of its unfolded effector domain region and the CPD across the plasma membrane. The eukaryotic host-specific molecule inositol hexakisphosphate activates CPD in the cell cytosol, inducing autoproteolytic cleavage at specific leucine residues between the arrayed effector domains (*28*). This mechanism promotes the release of multiple individual effectors, each targeting specific host cellular processes, including the disruption of actin assembly (*30–34*),the inactivation of small GTPases (*34–36*), and the inhibition of autophagic and endocytic trafficking (*5*, *8*). Together, the effector-induced cytopathic activities and the impairment of cell integrity through pore formation lead to host cell death.

Although the mechanisms for MARTX toxin secretion, autoprocessing, and the intracellular localization and functions of almost all its effectors are well understood, the specific mechanism by which MARTX toxins initially bind to host cells remains elusive. Prohibitin 1 (PHB1) was initially suggested as a putative host receptor for the *V. vulnificus* MARTX toxin based on the observation that PHB1 can bind to a specific effector of the MARTX toxin of *V. vulnificus* strain CMCP6 (*37*). The study showed that PHB1 localized to the plasma cell membrane during cell intoxication, portending a positive feedback mechanism to facilitate further toxin binding (*37*). However, it is critical to consider that PHB1, when present on the plasma membrane, predominantly localizes to the inner leaflet of the lipid bilayer (*38*), questioning whether PHB1 would be accessible at host surfaces ahead of pore formation and effector translocation. The lack of any other investigation into MARTX toxin surface interactions underscores the novelty and necessity of investigating MARTX toxin host receptors.

In this study, we aim to address the nature of the MARTX receptor given the broad spectrum of cells susceptible to this toxin. First, we identify a 273 aa region of the toxin that binds cells surfaces. This receptor binding domain has tandem immunoglobulin (Ig)-like folds, herein called Aknot. We then discover that the interactions of Aknot with host cells are governed by complex N-glycans on the ectodomain of L1 Cell Adhesion Molecule (L1CAM) and other select N-glycoproteins. We demonstrate that the binding of Aknot to N-glycans is governed by the inner obligate N-acetylglucosamine (GlcNAc) residues, a motif ubiquitous on cell surfaces, explaining the broad susceptibility of most cell types to these cytotoxins.

## RESULTS

### A MARTX toxin region with a predicted tandem Ig-like fold binds host cell surfaces

Since the CPD and the effector domain of MARTX_Vv_ are not required for receptor binding (*39*),we focused on identifying a surface receptor binding domain within the N- and C-terminal repeat regions of the MARTX_Vv_. Utilizing the AlphaFold2 (AF2) structure prediction algorithm (*40*), we scanned for structural distinct features within the extended conserved regions of the toxin that might suggest a binding module for interaction with mammalian cell surfaces. Overall, the algorithm predicted the tandem repeat regions with the greatest confidence compared to the linker region or the non-tandem repeat regions (fig. S1, B and C). AF2 predicted the “A” and “B” repeat regions as beta-roll structures, similar to prior analysis (Fig.1B and ref. (*7*)). As previously noted, the model for the tandem repeat regions predicted amphipathic features at the turns of the beta-roll, suggesting a potential role of these repeats for forming pores that span host membranes (fig. S1,D and E and refs.(*9*, *17*)).

The structural model also predicted with high confidence a 273-aa insert between the 10th and the 11th A repeats that extends away from the beta-roll and structurally appears like a two-headed rope knot, a region herein called the Aknot region (Fig.1B). The region was predicted to fold into two symmetrical subdomains, each comprised of a five-stranded beta sheet Ig-like fold interconnected by two antiparallel beta-strands (Fig. 1C). Since Ig-like folds are conducive to host surface interactions (*41*), we hypothesized that Aknot may engage host surface receptors. To test this hypothesis, the Aknot region of the MARTX toxin from *V. vulnificus* CMCP6 was recombinantly purified with an N-terminal 6xHis-Flag tag for affinity purification and a C-terminal hemagglutinin (HA) tag for antibody detection. This protein (Aknot-HA) was incubated with multiple cell types across a concentration gradient at 4°C to facilitate binding while inhibiting endocytosis. Following binding, cells were lysed for Western blot analysis to detect bound Aknot in whole cell extracts. In addition, live cells were subjected to flow cytometry after binding to quantify Aknot-HA on the surface. Furthermore, cellular immunofluorescence was performed to visualize surface-bound Aknot-HA on non-permeabilized cells. Both Western blotting (Fig.1D) and flow cytometry assays (Fig.1E) showed a concentration-dependent binding of Aknot-HA to HeLa epithelial cells and other commonly cultured cell lines (fig. S2, A to C) in the nanomolar concentration range. With immunofluorescence imaging, we demonstrated that Aknot serves as a binding domain for both T84 intestinal epithelial cells relevant to the food-borne route of *V. vulnificus* infection (Fig.1, F and G) and skin fibroblasts relevant to wound infection (fig. S2D).

### Aknot binds to L1CAM on the host surface

To identify eukaryotic membrane proteins that Aknot recognizes, we employed affinity purification mass spectrometry (AP-MS) to identify interacting partners of Aknot from the whole cell lysates. Mass spectrometry analysis of the isolated fractions from two independent experiments revealed peptides corresponding to various membrane proteins (Data S1). While the specific proteins identified varied between the two experiments, the only membrane protein consistently detected and ranked among the top 10 candidates by peptide coverage in both experiments was L1CAM, a single-pass membrane protein belonging to the Ig superfamily of cell adhesion molecules (Table S1). We used Western blotting to validate that Aknot-HA pulled down the L1CAM protein from HeLa cell lysates (fig. S3A).

To validate Aknot-HA and L1CAM binding in vitro, wells of a high adsorbent plate were coated with a gradient of the ectodomain of recombinant human L1CAM purified from human embryonic kidney 293 (HEK293) cells, and the wells were subsequently incubated with increasing concentrations of Aknot-HA. At ≥ 0.1 μM concentrations of Aknot, a binding curve was observed upon L1CAM titration, confirming the specific interaction between Aknot-HA and L1CAM (Fig.2A).

**Fig. 2.**
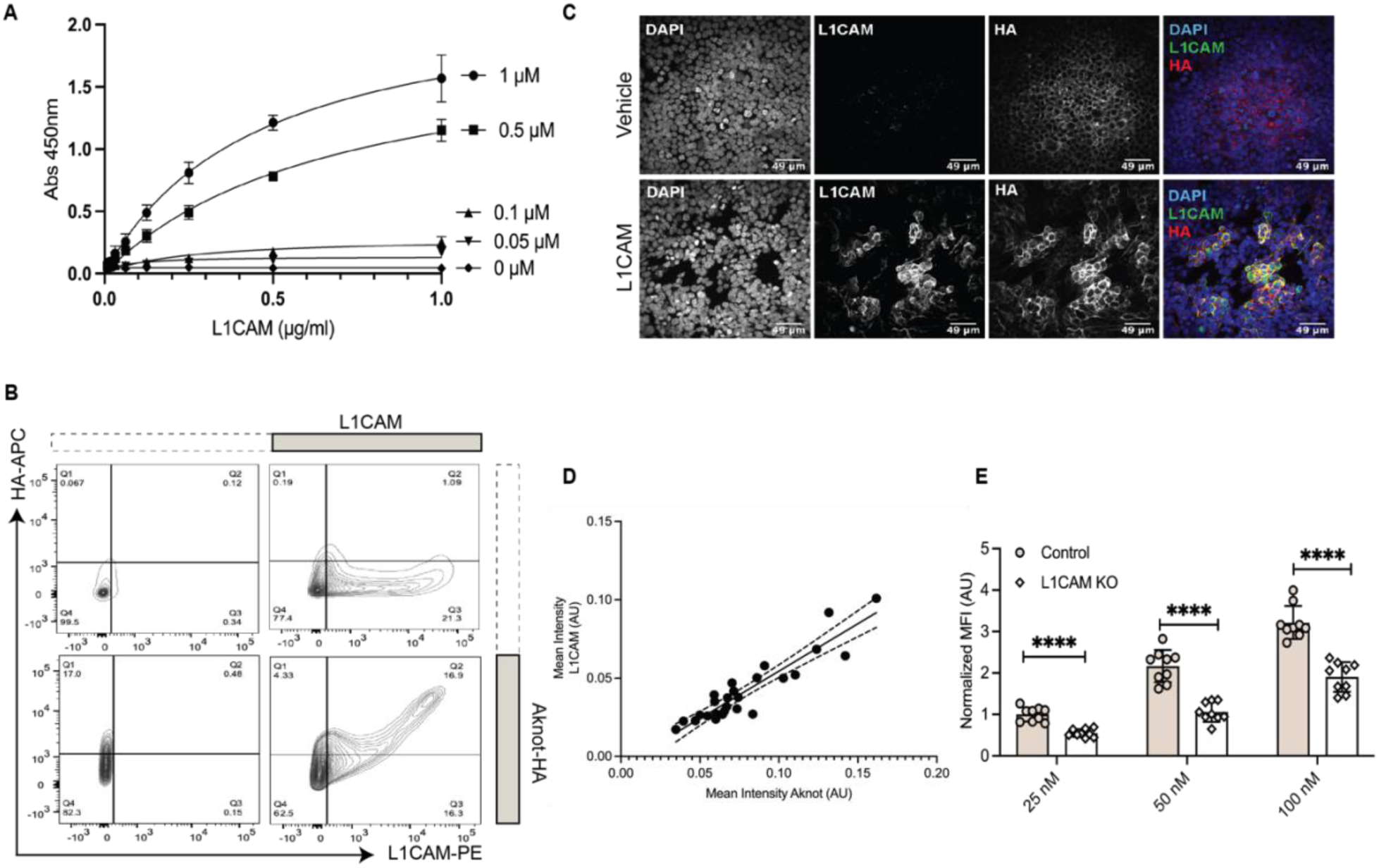
Aknot interacts with L1CAM. (A) ELISA binding curve of Aknot-HA to the ectodomain of L1CAM. High-absorbance plates were coated with L1CAM across a concentration gradient, reaching up to 1 μg/ml. The binding intensities Aknot-HA were assessed across a range of specified concentrations. Data points are presented as mean ± SD from *n=*3 individual experiments and fitted to a one-site specific binding least squares fit model in Prism 10. (B) Representative flow cytometry plots of Aknot-HA binding to HEK293T cells. Cells were transfected with either an empty vector (pcDNA3.1) or the vector with L1CAM coding sequence. After 48 hr, cells were incubated with 100 nM of Aknot-HA or PBS. The binding signal of Aknot-HA was detected by Alexa Fluor 647-conjugated anti-HA; surface expression of L1CAM was detected by PE-conjugated anti-L1CAM. (C) Immunofluorescence imaging of Aknot-HA and L1CAM on non-permeabilized HEK293T cells transfected with an empty vector (pcDNA3.1) or the vector with L1CAM coding sequence. After 48 hr, cells were incubated with 500 nM of Aknot-HA. (D) Correlation of L1CAM and Aknot-HA mean pixel intensity for each cell in panel C quantified from *n=*3 individual experiments analyzed by CellProfiler. Simple linear regression was fitted in Prism 10, best-fit was shown in solid line, 95% confidence intervals were shown in dashed line (R^2^ = 0.85). (E) Quantification of flow cytometry analysis of Aknot-HA binding to control and L1CAM^-/-^ HeLa Cas9 cells detected by Alexa 647-conjugated Anti-HA antibody. The average mean fluorescence intensity was normalized to 25 nM Aknot-HA binding to control cells for each experiment. Data are presented as mean ± SD from *n=*3 individual experiments. (\*\*\*\**p*<0.0001, Student’s *t* tests)

As our AP-MS data suggested that Aknot has other binding partners, we examined whether L1CAM is sufficient for Aknot binding. Since HEK293T cells naturally express deficient levels of L1CAM (fig.S3B), we incubated Aknot-HA with HEK293T cells with and without ectopic overexpression of full-length human L1CAM (Fig.2B). Using the flow cytometry assay, we detected a low level of binding of Aknot-HA to HEK293T cells transfected with the vehicle vector. In contrast, the binding of Aknot-HA to HEK293T cells increased in correlation with the overexpression of L1CAM on the surface of HEK293T cells (fig. S3C). Consistently, using confocal imaging, we observed that Aknot-HA bound preferentially to HEK293T cells overexpressing L1CAM (Fig.2, C and D). These results indicate that L1CAM can sufficiently recruit Aknot to the host surface to enhance binding.

We next determined the extent to which L1CAM is required as a receptor for Aknot using HeLa cells, which naturally express high levels of L1CAM (fig. S3B). A CRISPR Cas9-based system (*42*) was used to generate isogenic *L1CAM*^-/-^ cells with single-cell cloning (fig.S3, C and D). Compared to control cells introduced with scrambled small guide RNA, *L1CAM^-/-^* cells showed reduced binding of Aknot, exhibiting 44 ± 13% reduction of binding at 25 nM, 51 ± 7% reduction at 50 nM, and 40 ± 12% reduction at 100 nM (Fig.2E). The elimination of L1CAM on HeLa cell surfaces, while failing to abolish Aknot binding, reduced Aknot binding by approximately half.

Together, the notable increase of Aknot binding induced by L1CAM and the decrease in binding in the absence of L1CAM, suggest that L1CAM is a major receptor for Aknot. However, as residual binding to *L1CAM*^-/-^ cells persisted, L1CAM must not be the only receptor involved in Aknot binding on the host surface.

### Aknot targets complex N-glycans

The surface exposed L1CAM ectodomain is highly N-glycosylated, carrying over 20 N-glycosylation sites (*43*). To probe the potential involvement of these N-glycans on L1CAM in Aknot-HA binding, we tested *in vitro* binding of Aknot-HA to the ectodomain of L1CAM with and without N-glycans by use of the peptide-N-glycosidase F (PNGaseF), an endoglycosidase that cleaves and releases all N-glycans from glycoproteins. In the absence of its N-glycans, Aknot-HA failed to bind L1CAM, indicating that the N-glycans are essential for this interaction (Fig. 3A).

**Fig. 3.**
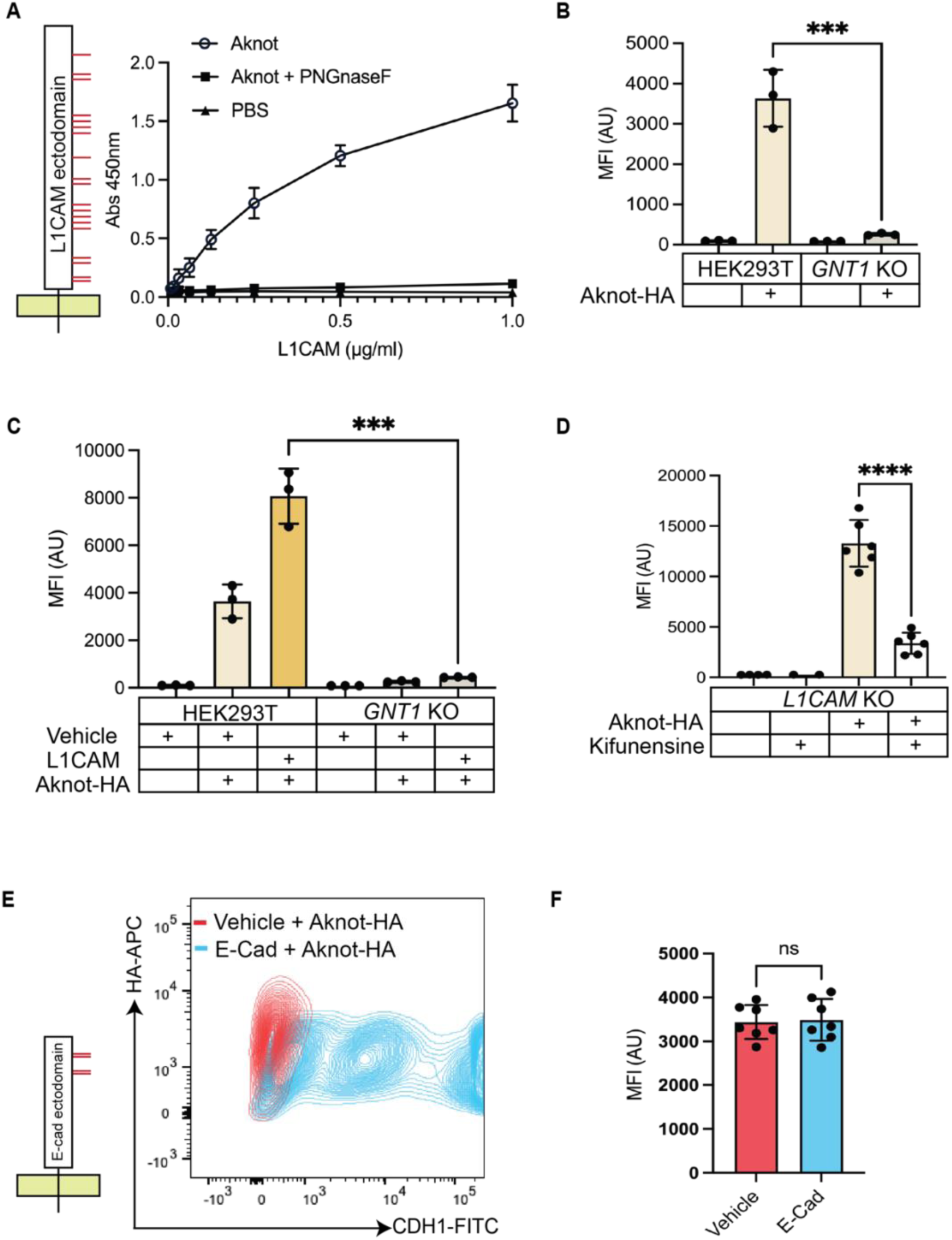
Aknot binds to host surface complex N-glycans. (A) ELISA binding curve of Aknot-HA to the ectodomain of L1CAM treated with and without PNGnaseF. High-absorbance plates were coated with L1CAM across a concentration gradient, reaching 1 μg/ml, and treated with and without PNGnaseF at 37°C overnight. The binding intensities of Aknot-HA were assessed at 1 µM. Data points are presented as mean ± SD from *n=*3 individual experiments fitted to a one-site specific binding least squares fit model in Prism 10. The N-glycosylation sites on the L1CAM ectodomain are indicated as red straight lines, with the yellow box indicating cell membrane. (B) Flow cytometry analysis of 100 nM of Aknot-HA binding to HEK293T and *GNT1*^-/-^cells. Data points are presented as mean ± SD from *n=*3 individual experiments (\*\*\**p*<0.001, student’s *t* test). (C) Flow cytometry analysis of Aknot-HA binding to HEK293T and *GNT1*^-/-^cells. Both cell lines were transfected with either an empty vector (pcDNA3.1) or the vector encoding the L1CAM sequence. 48 hours after transfections, cells were incubated with 100 nM of Aknot-HA or PBS. The binding signal of Aknot-HA was detected by Alexa Fluor 647-conjugated anti-HA. Data points are presented as mean ± SD from *n=*3 individual experiments (\*\*\**p*<0.001, Student’s *t* test). (D) Flow cytometry analysis of 100 nM of Aknot-HA binding to L1CAM^-/-^ HeLa Cas9 cells treated with or without kifunensine at 37°C overnight. Data points are presented as mean ± SD from *n=*3 individual experiments (\*\*\*\**p*<0.0001, Student’s *t* test). (E) Representative flow cytometry plots of Aknot-HA binding to HEK293T cells. Cells were transfected with either an empty vector (pcDNA3.1) or the vector encoding the GFP-tagged E-cadherin sequence (CDH1-GFP). 48 hours after transfections, cells were incubated with 100 nM of Aknot-HA. The binding signal of Aknot-HA was detected by Alexa Fluor 647-conjugated anti-HA. The N-glycosylation sites on the E-cadherin ectodomain are indicated as red straight lines, with the yellow box indicating cell membrane. (F) Quantifications of Aknot-HA binding in panel E on cells transfected with the vehicle vector and cells successfully transfected with the vector encoding E-cadherin. Data were represented as mean ± SD from *n=*3 individual experiments (ns, not significant, Student’s *t* test).

To dissect the role of N-glycans in Aknot-HA binding, we tested whether complex N-glycans with elaborate glycan structures were required by employing HEK293T cells genetically engineered to be deficient in biosynthesis of complex N-glycans (HEK293T *GNT1^-/-^*) by knockout of the *GNT1* gene encoding the first decisive step in conversion of oligomannosyl N-glycans to complex type. HEK293T *GNT1*^-/-^ cells only express oligomannosylated N-glycans, and Aknot-HA did not bind to these (Fig. 3B), even when L1CAM was ectopically overexpressed (Fig.3C). These data demonstrate that the increased Aknot binding observed in L1CAM-expressing HEK293T cells resulted at least in part from complex N-glycans on L1CAM. The residual binding of Aknot-HA to HeLa *L1CAM*^-/-^ cells (Fig. 2D) may result from Aknot binding to complex N-glycans on other glycoproteins. To confirm this finding, we pretreated HeLa *L1CAM*^-/-^ cells with kifunensine, a mannosidase inhibitor that prevents complex N-glycan biosynthesis, before binding with Aknot-HA (Fig.3D). As predicted, the drug treatment resulted in a 73 ± 9% reduction in Aknot-HA binding to HeLa *L1CAM*^-/-^, suggesting that Aknot recognizes the complex N-glycans on host membrane N-glycoproteins other than L1CAM.

Next, to explore the requirement of Aknot-HA binding to particular carrier proteins with complex N-glycans, we overexpressed E-cadherin (CDH1), a well-studied complex-type N-glycoprotein that functions in cell adhesion similar to L1CAM. Notably, Aknot-HA binding to HEK293T cells was not increased when CDH1 was introduced (Fig. 3, E and F). This demonstrates that Aknot exhibits a degree of selectivity towards N-glycoproteins, suggesting that structures and spacing of multiple complex N-glycans, likely well represented by L1CAM, are involved in the recognition.

### Aknot binds preferentially to inner GlcNAc residues in biantennary N-glycans

To further investigate Aknot-HA binding to distinct structural features of complex N-glycans, we requested a screening of Aknot-HA from the Consortium for Functional Glycomics (CFG) mammalian-type glycan microarrays containing 562 oligosaccharides. However, Aknot-HA did not produce significant binding intensities to any tested N-glycans (Data S2). This was consistent with our finding that Aknot recognizes particular presentations of multiple N-glycan motifs on select glycoproteins. We therefore turned our efforts towards cell-based glycan arrays that enable display and structural dissection of N-glycan binding in the context of proteins and the cell membrane (*44*). Cell-based glycan arrays take advantage of the stepwise biosynthesis of most types of cellular glycosylation (Fig. 4A) and employ genetic engineering (KO/KI) of key glycosyltransferase genes in human HEK293 cells to generate cell libraries with loss/gain of particular structural glycan features (*45*). These libraries of isogenic cells display N-glycoproteins with distinct N-glycan structures on the cell surface for interrogation by flow cytometry. To investigate the glycoconjugate types involved in Aknot binding, we tested the interaction between Aknot-HA and a panel of HEK293 cells lacking O-glycans, glycolipids, or complex N-glycans, as well as cells with combinations of these glycoconjugate types (Fig. 4B). Aknot-HA binding was only affected by loss of complex N-glycans (HEK293KO:MGAT1), which indicates that Aknot binds specific features found on N-glycans that are not shared with glycolipids and O-glycans, such as N-acetyllactosamine (LacNAc) and sialic acid capping epitopes that are often recognized by microbial adhesins (*46*, *47*). Next, we tested Aknot-HA binding to a cell library designed to probe the required branching state of N-glycans (Fig. 4C). This revealed that Aknot-HA binding was significantly reduced when the synthesis of biantennary N-glycans was abrogated (HEK293KO:MGAT2), while interfering with tri- and tetraantennary N-glycans had only minor effects. Note that while individual targeting of the bisecting N-glycosylation step (HEK293KO:MGAT3) did appear to reduce Aknot-HA binding, the combinatorial targeting of bisecting and tri/tetraantennary N-glycosylation (HEK293KO:MGAT3/4A/4B/5) did not, indicating that the critical structures required for Aknot-HA binding are biantennary N-glycans.

**Fig. 4.**
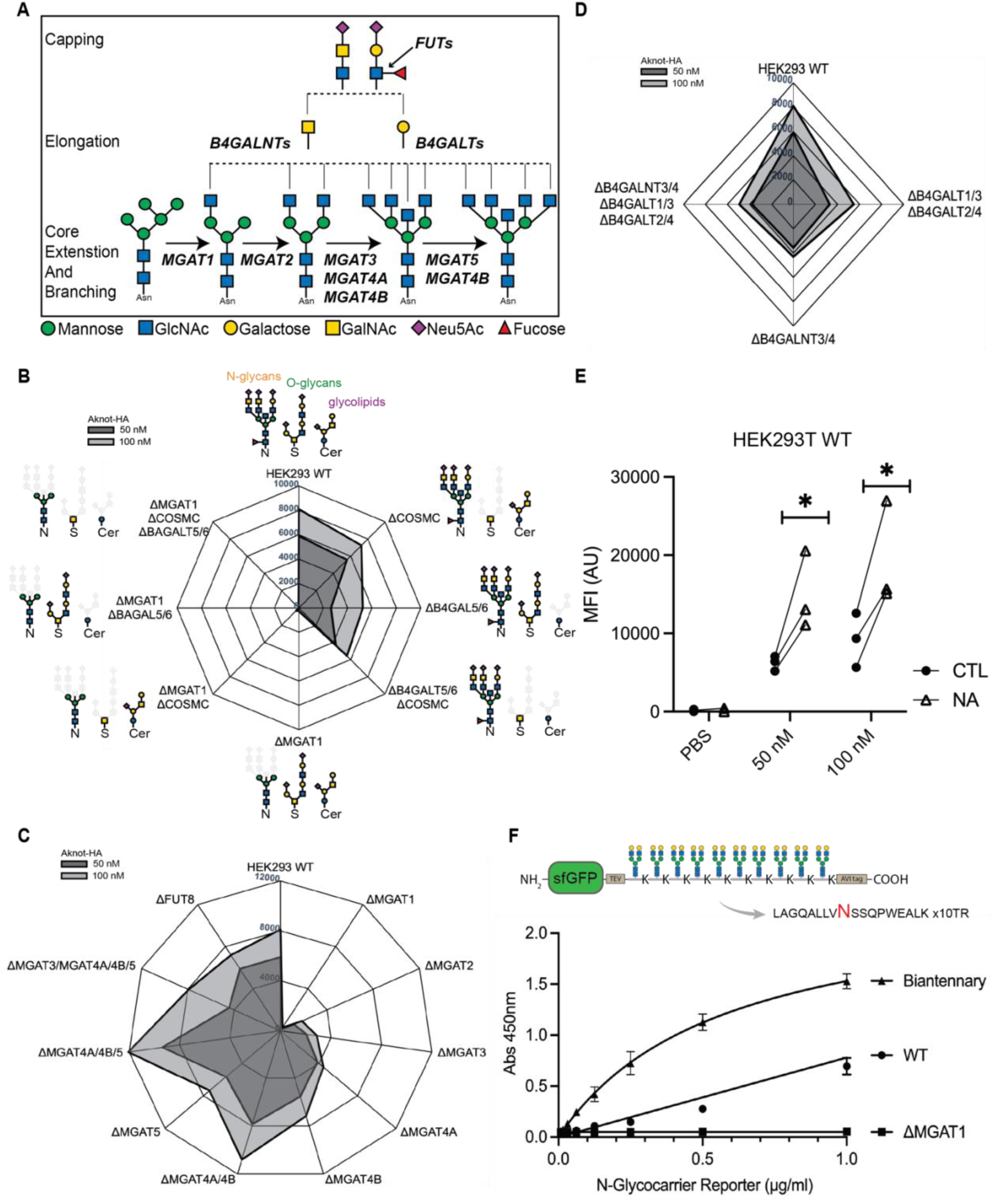
Aknot binds to GlcNAc on complex N-glycans. (A) Schematic representation of key glycosyltransferase genes required in complex N-glycan synthesis pathways. (B) Radar plot illustrating the mean MFI of Aknot-HA binding at 50 nM and 100 nM to genetically engineered HEK293 cells with specific genes knocked out in the synthesis pathways of O-glycans (HEK293KO:COSMC), glycolipids (HEK293KO:B4GALT5/6), and N-glycans (HEK293KO:MGAT1), as determined by flow cytometry analysis. Each data point represents the mean MFI from *n*=3 individual experiments. (C) Radar plot illustrating the mean MFI of Aknot-HA binding at 50 nM and 100 nM to genetically engineered HEK293 cells with specific genes knocked out in the N-glycan core branching pathways (HEK293KO:MGAT2, HEK293KO:MGAT3, HEK293KO:MGAT4A, HEK293KO:MGAT4B), as determined by flow cytometry analysis. Each data point represents the mean MFI from *n*=3 individual experiments. (D) Radar plot illustrating the mean MFI of Aknot-HA binding at 50 nM and 100 nM to genetically engineered HEK293 cells with specific genes knocked out in the N-glycan elongation pathways that result in the absence of LacDiNAc (HEK293KO:B4GALNT3/4) and LacNAc (HEK293KO:B4GALT1/2/3/4), as determined by flow cytometry analysis. Each data point represents the mean MFI from *n*=3 individual experiments. (E) Flow cytometry analysis of Aknot-HA binding to wild-type HEK293 cells pre-treated with or without neuraminidase at 37°C for 1 hour. Data points were plotted from *n=*3 individual experiments, with points from the same experiment connected (\**p*<0.01, Student’s *t* test). (F) ELISA binding curve of Aknot-HA to artificial purified N-Glycocarrier reporters containing 10 N-glycans and expressed in glycoengineered CHO cells as indicated in the cartoon illustration. Data points are presented as mean ± SD from *n=*3 individual experiments and fitted to a one-site specific binding least squares fit model in Prism 10.

These findings suggest that Aknot-HA does not recognize simple terminal glycan epitopes. To substantiate this, we employed a cell library with deficiencies in the basic terminal core structures LacNAc (Galβ1-4-GlcNAcβ1-R) and LacDiNAc (GalNAcβ1-4GlcNAcβ1-R) (Fig. 4D). Aknot-HA binding was unaffected by loss of these extension of the core GlcNAc residues on N-glycans. In addition, neuraminidase treatment, which enzymatically removed terminal sialic acids on N-glycans, did not abolish but enhanced Aknot-HA binding (Fig. 4E). These results indicate that Aknot-HA recognizes the GlcNAc residues on biantennary N-glycans independent of further galactosylation and sialylation modifications. This interpretation was supported by the finding that Aknot-HA binding to L1CAM in ELISA assays was inhibited by high concentrations (>100μM) of the trisaccharide GlcNAc_3_ (GlcNAcβ1-4 GlcNAcβ1-4GlcNAc), while up to 1 mM of the GlcNAc monosaccharide had nearly no effect (fig. S4A).

To further confirm that Aknot binds N-glycoproteins with presentation of multiple N-glycans, we employed a recombinant artificial N-Glycocarrier reporter designed with 10 definably spaced N-glycans (*48*). We expressed the N-Glycocarrier reporter in genetically engineered CHO cells to produce defined N-glycan variants and used the isolated reporters to evaluate Aknot-HA binding by ELISA (Fig. 4F). Aknot-HA bound to the reporter with the mixture of bi- to tetraantennary complex sialylated N-glycans produced by CHO wild-type cells, while no binding to the reporter with homogenous high-mannose N-glycans (CHO KO:MGAT1) was detected. In contrast, when probing the reporter with homogenous biantennary complex N-glycans without sialic acid, binding was increased compared to wild-type. This confirms that Aknot binding requires multiple biantennary N-glycans with specific spatial presentations and that such binding motifs can be presented by an artificial protein backbone.

### MARTX_Vv_ Aknot region is essential for *V.vulnificus* virulence

Having identified that Aknot binds to the host surface via specified motifs on complex N-glycans, it is essential to determine the function of this 273-aa Aknot region within the context of the larger 5206-aa MARTX_Vv_. To assess how Aknot fulfills its receptor-binding role in promoting MARTX_Vv_ intoxication during *V.vulnificus* infections, we first evaluated the significance its receptor, complex N-glycans, as the receptor for the holotoxin. Since no group has successfully purified a full-length active MARTX toxin for *in vitro* or *in vivo* studies, we determined host susceptibility mediated by cell surface complex N-glycans to MARTX_Vv_ by incubating HEK293T wild-type and *GNT1*^-/-^ cells with live *V. vulnificus* that actively export MARTX_Vv._ This was followed by lactate dehydrogenase (LDH) release assays to detect host cell lysis induced by toxin pore formation. To eliminate confounding effects caused by hemolysin, an additive cytotoxin of *V.vulnificus*, we disrupted the hemolysin-encoding *vvhA* gene on the bacterial chromosome in situ (CMCP6*ΔvvhA*). Additionally, since the effectors of the MARTX toxin do not play a role in host surface interactions, we used a variant of *V. vulnificus* in which the effector domain sequence within the *rtxA1* gene encoding MARTX was replaced with the sequence encoding beta-lactamase (CMCP6*ΔvvhA rtxA1:bla)* (Fig. 5A). Co-culturing cells with this effector-free variant strain (CMCP6*ΔvvhA rtxA1:bla)* led to cell death within 2 hours, however, no differences in host susceptibility to toxins were detected between HEK293T wild-type and *GNT1^-/-^* cells (Fig. 5B). These results demonstrate that in the absence of Aknot binding partners on the host, *V. vulnificus* delivered MARTX toxin can still mediate sufficient surface interaction to drive pore formation. This further indicates that there must be other receptors in addition to complex N-glycans, requiring additional receptor-binding domains besides the Aknot region.

**Fig. 5.**
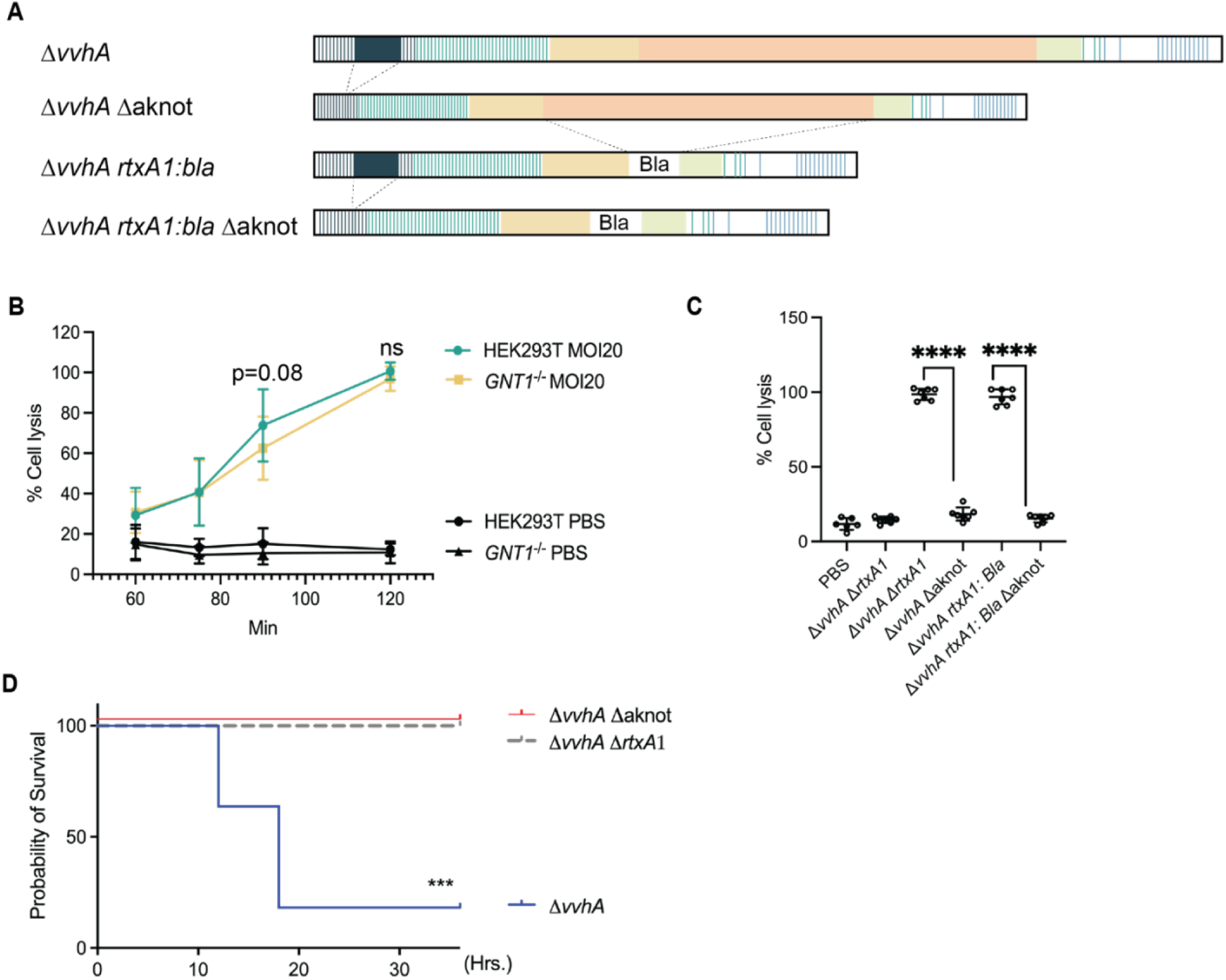
MARTX_Vv_ has additional receptor-binding domain targeting different receptors. (A) Schematic representation of the structure organizations of the MARTX toxin variants. (B) Temporal analysis of cell lysis induced by *V. vulnificus* CMCP6 Δ*vvhA* effector-free strain (*Δvvha rtxA1:bla*) in HEK293T and *GNT1*^-/-^ cells at MOI=20. Lysis was measured by lactate dehydrogenase (LDH) activity in the culture medium at 30-minute intervals for 2 hours. Data points are presented as mean ± SD from *n=*3 individual experiments (ns; not significant, Student’s *t* test). (C) Cell lysis of HEK293T cells intoxicated by indicated strains at MOI=20 after 180 mins measured by LDH. Data points are presented as mean ± SD from *n=*3 individual experiments (\*\*\*\**p*<0.0001, Student’s *t* test). (D) Survival curve of mice (*n*=10 for each condition) intragastrically infected with 4 x10^7^ colony-forming units of the indicated strain and monitored for 36 hours. Survival differences assessed by Kaplan-Meier analysis were statistically compared using the log-rank test.

Since the Aknot region might not be the sole MARTX toxin receptor-binding domain, we evaluated the significance of Aknot itself during intoxication by co-culturing HEK293T cells with variants of *V. vulnificus* in which the *rtxA1* codons for Aknot were removed in-frame(Δ*aknot*) (Fig. 5C). Mutated strains with Aknot removed from the toxin failed lyse cells, similar to the toxin-null strain (Δ*rtxA1*), indicating that the absence of Aknot region rendered the toxin incapable of forming pores in cell membranes. To recapitulate this essential role of Aknot *in vivo*, mice were infected with *V. vulnificus* strain variants following a food-borne infection model. A lethal dose of *V. vulnificus* strain CMCP6Δ*vvhA* that secretes a MARTX toxin that retains all of the effector domains succumbed to the intestinal infection within 36 hours. By comparison, mice infected with Δ*aknot* variant all survived, mimicking the phenotype of mice infected with the toxin null strain Δ*rtxA1*. Thus, a MARTX toxin variant that lacks the Aknot region is entirely avirulent during lethal intestinal infection. These data support that Aknot is indispensable for MARTX-dependent intoxication during mammalian intestinal infection.

## DISCUSSION

The role of the MARTX toxin as a pivotal virulence factor of *V. vulnificus* has been extensively studied, with prior research predominantly focused on the functions of its enzymatic effectors. This study unravels the mechanism through which the toxin initiates its intoxication at the host cell surface. We discover the Aknot module as a receptor binding domain that engages N-glycans on select glycoproteins. The recognized motif is unusual among toxin receptors as it is comprised of the internal obligate GlcNAc residues of complex biantennary N-glycans rather than the terminal sugars. Targeting such internal motifs in glycans provides broader interactions with glycoproteins independent of variable modifications of terminal glycans, which are often partly degraded by microbial glycoside hydrolases. Moreover, we provide evidence that Aknot exhibits selectivity for particular N-glycoproteins such as L1CAM, which we predict is guided by patterns of multiple N-glycans that can be arrayed when found on proteins densely covered by N-glycans, such as L1CAM.

We showed that L1CAM is an important host receptor for Aknot, as its high expression correlates with increased Aknot binding. In the context of *V. vulnificus* food-borne infections, L1CAM is likely to serve as a biologically relevant receptor because it is well expressed in the endothelial cells of the human gastrointestinal tract and human dermal tissues (*49*). Notably, the expression of L1CAM is significantly elevated when the integrity of the intestinal epithelium is disrupted (*50*), a condition that occurs during *V. vulnificus* infections. Our observed L1CAM-dependent increase in Aknot binding to cell surfaces suggests the *V. vulnificus* MARTX toxin could capitalize on this host response.

Importantly, our data also showed that Aknot binds to L1CAM only when it is N-glycosylated, adding MARTX toxin to the expanding list of bacterial toxins that target mammalian glycans as host receptors. However, most if not all the other glycan-binding toxins recognize terminal glycan moieties, since the recognition of specific terminal glycan modifications allows toxins to bind to and enter only highly specific cell types (49–53). By contrast, Aknot distinctively engages a N-glycan binding motif that comprises of multiple clustered biantennary N-glycans, at least partly through recognition of the obligate internal GlcNAc residues which are the fundamental components of all complex N-glycans. Notably, the biantennary N-glycan structure is also the most abundant structural class across various tissues (*55*). This ubiquity significantly expands the host range for Aknot binding, explaining the host tropism of MARTX_Vv._ Furthermore, Aknot exhibits an unusual binding specificity towards a plurality of N-glycans. To our knowledge, only the broadly neutralizing anti-HIV antibody 2G12 has been demonstrated to engage a glycan epitope comprised of multiple N-glycans (*56*). We demonstrated strong binding of Aknot to artificial N-Glycocarrier reporters with evenly spaced N-glycans, which can now serve as a flexible scaffold to further interrogate the molecular nature of the Aknot binding epitope and provide access to defined ligands for structural studies of the binding mode.

Our results demonstrate that the host receptor for Aknot is important but not essential for surface binding by the holotoxin. This suggests a redundancy of host receptors during MARTX_Vv_ intoxication involving multiple receptor-binding domains, such that the reduction in intoxication efficiency due to the loss of one class of receptors is not sufficient to abolish toxin translocation. Targeting multiple receptors on host surface represents a widespread mechanism bacterial toxins employ to enhance their virulence, expand the spectrum of susceptible hosts, and avoid cell resistance by downregulation of a single receptor. For example, *Clostridioides difficile* Toxin B (TcdB) targets a suite of distinct receptors via discrete receptor-binding domains, providing a diverse physiological context for infection. These receptors include Frizzled 1, 2 and 7 (FZD1/2/7), chondroitin sulfate proteoglycan 4 (CSPG4); nectin-3/poliovirus receptor-related protein 3 (PVRL3); and tissue factor pathway inhibitor (TFPI). Notably, even the combined genetic knock-out of *FZD1/2/7* and *CSPG4* did not result in cell resistance to TcdB intoxication (*57*). Determining other receptor-binding domains on MARTX_Vv_ in future studies will deepen our understanding of MARTX toxin host range and mechanism of infection. A reduction or elimination of host susceptibility towards MARTX_Vv_ will likely be achieved once more receptors are removed from the host surface. Additionally, the functional significance of Aknot beyond receptor-binding should be further investigated, as our results demonstrated that Aknot is required for *in vivo* infection, likely by facilitating multiple stages in the process of MARTX_Vv_ entry into host cells.

In conclusion, the findings presented here pave the way for understanding the mechanisms by which MARTX toxins interact with the host cell surface by identifying a receptor-binding domain on MARTX_Vv_ and establishing N-glycan motifs on select N-glycoproteins as its receptors. A better understanding of residues on the Aknot that are crucial for N-glycan binding is an essential next step in understanding the role of N-glycans as receptors for MARTX_Vv_ across different *V. vulnificus* strains and other *Vibrio* species. If these residues are conserved across species, it would strongly suggest that N-glycans serve as universal receptors for MARTX toxins, indicating a shared strategy for host interaction and infection among different strains. If those residues are not conserved, it suggests the diversity of receptors utilized by toxins from various species, providing insights into host specificity and the evolutionary trajectory of these pathogens.

## MATERIALS AND METHODS

### Protein expression and purification

The construct encoding Aknot-HA (residues 244– 571 of the MARTX toxin from *V. vulnificus* CMCP6), with a 6xHis tag and a C-terminal HA tag or with an N-terminal Avidin (Avi) tag (Avi-Aknot), was cloned into the pET28A+ vector by Twist Bioscience (South San Francisco, CA). The plasmid was transformed into *Escherichia coli* BL21(DE3) Star cells. Cultures were grown at 37°C in Terrific Broth (TB) medium until the optical density at 600 nm reached 1.0. Protein expression was then induced with 0.5 mM isopropyl β-d-1-thiogalactopyranoside (IPTG) and cultured overnight at 25°C. Cells were harvested by centrifugation at 6,000 x *g* for 10 minutes and resuspended in lysis buffer containing 10 mM Tris-HCl (pH 8.3), 500 mM NaCl, 10% glycerol, 0.1% IGEPAL CA-630, and cOmplete cocktail inhibitor (Roche). Cells were then sonicated for 1 hour, followed by clarification of the lysate by centrifugation at 38,759 xg for 40 minutes at 4°C. Protein purification was performed using a HisTrap FF column (Cytiva, Catalog #17525501) on an ÄKTA Purifier system. The column was equilibrated with binding buffer (10 mM Tris-HCl, pH 8.3, 500 mM NaCl, 10 mM imidazole). After loading the lysate, the column was washed with high-salt buffer (10 mM Tris-HCl, pH 8.3, 1 M NaCl, 25 mM imidazole), followed by elution of the target protein with elution buffer (10 mM Tris-HCl, pH 8.3, 500 mM NaCl, 500 mM imidazole). Eluted fractions were dialyzed against 10 mM Tris-HCl (pH 8.3), 500 mM NaCl to remove imidazole and then further purified by size-exclusion chromatography using HiLoad^TM^ 26/600 Superdex^TM^ 200 pg column (Cytiva, Catalog #28989336) pre-equilibrated with the same buffer. Fractions containing the target protein, as determined by SDS-PAGE analysis, were pooled and concentrated using a 10-kDa molecular weight cutoff concentrator. The final protein solution was adjusted to include 20% glycerol, flash-frozen in liquid nitrogen, and stored at -80°C for future analyses.

### Cell culture

HeLa cells, including genetically engineered variants, were maintained in Dulbecco’s Modified Eagle Medium (DMEM) (Gibco, Catalog #11965-092) supplemented with 10% fetal bovine serum (FBS) and 100 U/ml penicillin-streptomycin. HEK293T cells, HEK293T GNT1^-/-^ (ATCC, Catalog #CRL-3022), HEK cells, and their engineered counterparts were cultured in a 1:1 mixture of Ham’s F12 medium and DMEM (F12/DMEM) enhanced with GlutaMAX™ (Gibco, Cat.#10565-018), supplemented with 10% FBS and 100 U/ml penicillin-streptomycin. All cell lines were incubated under a humidified atmosphere at 37°C with 5% CO_2_.

### Toxin binding

Indicated cells were seeded in 12-well plates and incubated overnight to achieve approximately 90% confluency by the following day. Toxin fragments were then incubated with cells in either fresh media supplemented with serum or in Hank’s Balanced Salt Solution (HBSS) (Thermo Scientific, Catalog #14025092) at the specified concentrations. This incubation occurred for 1 hour at 4°C on a rocker. Subsequently, cells were washed three times with phosphate-buffered saline (PBS).

### Western blots

Cells from each well of the 12-well plates were lysed using RIPA buffer (Thermo Scientific, Cat.# 89900) supplemented with protease inhibitor cocktails (Roche, Cat.#11836170001). The lysates were then clarified by centrifugation at 13,000 x *g* for 15 minutes. A third of the total volume of each supernatant were boiled and run on 4 to 12% Tris/Glysine polyacrylamide gels (BioRad, Cat.#456-1093). Following electrophoresis, proteins were transferred onto nitrocellulose or onto PVDF membranes for immunoblotting. The membranes were blocked and then incubated overnight at 4°C with the primary antibodies: Tubulin (Sigma, Cat.#T6074; Cell signaling, Cat.#2144), HA-tag (Sigma-Aldrich, Cat.#H6908), L1CAM (Santa Cruz, Cat.#SC-53386), Avidin (GenScript, Cat.#A00674). Membranes were washed and incubated with fluorescently labeled secondary antibodies: Goat anti-rabbit IgG conjugated with a 680 nm dye (LI-COR Biosciences, Cat.# 926-68071) and Goat anti-mouse IgG conjugated with an 800 nm dye (LI-COR Biosciences, Cat.# 926-32210), for 1 hour at room temperature. Proteins were detected under the LI-COR Odyssey M imaging system.

### Flow cytometry

Following toxin exposure, cells were washed and gently detached from each well using a FACS detaching buffer (25 mM HEPES pH 7.0, 2 mM EDTA, 10% bovine serum albumin (BSA)). The detached cells were incubated with an Alexa Fluor 647-conjucated anti-HA tag antibody (R&D, Cat.#IC6875R) at dilution of 1:20 in 100 µL of PBS containing 1% BSA (PBA) for 45 minutes in the dark at room temperature. Cells were then washed twice with PBA and analyzed using a BD FACSCanto II flow cytometer at Interdepartmental Immuno-Biology Flow Cytometry Core Facility of Northwestern University. 10,000 events were recorded for each sample and data was analyzed using FlowJo software.

For cell-based glycan array screening, a library of isogenic HEK293 cells with genetic engineered glycosylation capacities was used as previously described.(*44*) After toxin exposure, cells were incubated with APC conjugated anti-Flag tag antibody (Biolegend, Cat.#637308) at dilution of 1:1000 in 50 ul of PBS containing 1% BSA for 1hr in the dark. Cells were then washed twice with PBA and analyzed using Fortessa 3 at Flow Cytometry Core Facility of University of Copenhagen.

### Immunofluorescence imaging

Round glass coverslips (Ted Pella, Cat.#26022) were placed into each well of a 12-well plate. For HEK293T and HEK293T GNT1^-/-^ cells, these coverslips were pre-coated with poly-L-lysine (Advanced Biomatrix, Cat.#5048). Cells were seeded onto the coverslips and allowed to adhere overnight, followed by incubation with the specified toxin as previously described. Post-incubation, cells were washed with PBS and fixed with 4% paraformaldehyde for 20 minutes at room temperature. Cells were blocked in PBS containing 1% BSA, 10% normal goat serum, 0.3M glycine and 0.1% tween, for 1 hour at room temperature. This was followed by incubation with primary antibodies overnight at 4°C: HA tag(Cell Signaling, Cat.# 3724) and L1CAM(Abcam, Cat.# ab2435) diluted in PBS with 0.1% Tween-20 (PBST). After primary antibody incubation, cells were washed three times with PBST. For detection, coverslips were incubated with secondary antibodies: Goat anti-mouse Alexa Fluor 488 (Life Technologies, Cat.#A11001) and Goat anti-rabbit Alexa Fluor 647 (Life Technologies, Cat.#A21244) and DAPI (Invitrogen, Cat.#R37606) in PBST with 1% BSA in the dark at room temperature for 1 hour. Following a series of washes, coverslips were mounted onto glass slides using Prolong Gold antifade mountant (Life Technologies, Cat.#P10144). Imaging was performed at the Northwestern University Center for Advanced Microscopy using the Nikon A1R+ GaAsP Confocal Laser Microscope, and the acquired images were analyzed quantitatively using CellProfiler software.

### Cell transfection

HEK293T and HEK293T GNT1^-/-^ cells, at a density of 0.1 × 10^6^ cells, were seeded onto 12-well plates pre-coated with poly-L-lysine (Advanced Biomatrix, Cat.#5048) overnight. For transfection, 2.5 µg of plasmid DNA containing the *L1CAM* gene (Addgene, Cat.#89411), *CDH1-GFP* (Addgene, Cat.#28009) or the corresponding empty vector control pcDNA3 was prepared. Each plasmid was mixed with Lipofectamine 2000 (Invitrogen, Cat.#11668019) according to the manufacturer’s protocol for a 12-well plate format. Fresh media was added after 24 hours and cells were assayed at 48 hours post-transfection.

### Artificial N-Glycocarrier reporters

CHO cells with genetically engineered glycosylation capacities were used to produce an artificial N-Glycocarrier with 10 evenly spaced N-glycans. The N-Glycocarrier contains N-terminal sfGFP followed by a glycomodule comprised of 10 repeat 18-mer peptide N-glycosylation motifs (LAGQALLV**<u>N</u>**SSQPWEALK) and His purification tags. When expressed as a secreted fusion protein in glycoengineered CHO cells, the N-Glycocarrier obtains stoichiometric attachment of 10 N-glycans with structures that faithfully follow engineered glycosylation capacities.(*58*) Secreted N-Glycocarriers were isolated by Ni-NTA (ThermoFisher Scientific, Cat.#88221) chromatography and their glycans characterized by LC-MS/MS analysis of the trypsin released 18-mer glycopeptide.

### ELISA

MaxiSorp plates (Thermo Scientific, Cat.# 44-2404-21) were coated with the purified ectodomain of L1CAM (Sino Biological, Cat.#10140-H08H-100) or the respective recombinant N-Glycocarrier reporters in coating buffer (15 mM Na_2_CO_3_, 35 mM NaHCO_3_, pH 9.6) for 2 hours in 37°C. Wells were then blocked with PLI-P buffer (0.5 M NaCl, 3 mM KCl, 1.5 mM KH_2_PO_4_, 6.5 mM Na_2_HPO_4_,2H_2_O, 1% BSA, 1% Triton-X-100) for 1 hour at room temperature. Subsequently, indicated concentrations of toxin fragments were added to the wells and incubated for 1 hour at room temperature to allow for binding. Following the incubation period, the wells were washed with PBST and incubated with HRP-conjugated Anti-HA tag antibody at 0.5 µg/ml (GenScript, Cat.#A01296) for 1 hour at room temperature. The plates were then developed using TMB substrate (Thermo Fisher Scientific, Cat.# 34028) in the dark for 10 minutes. The enzymatic reaction was halted by adding quenching solution (0.5M H_2_SO_4_). The absorbance at 450 nm was measured using the microplate reader from Tecan Life Sciences.

For the GlcNAc competition assay, Aknot-HA was incubated with a titration of GlcNAc (Vector Laboratories, Cat.#S-9002) and GlcNAc_3_ (Neogen Corporation, Cat.#O-CHI3) in PLI-P buffer at room temperature for 1 hour. Subsequently, the Aknot-HA samples were applied to MaxiSorp plates coated with L1CAM and the ELISA procedure was performed as described above.

### PNGnaseF and Kifunesine treatment

Kifunensine (R&D Systems, Cat.# 3207) was applied to cultured cells at 0.02 mM in Opti-MEMTM reduced serum media (ThermoFisher, Cat.# 31985062) at 37°C overnight. PNGase F (Sigma-Aldrich, Cat.# 11365193001) was applied to ELISA plate at 0.5 U/well in 37°C overnight.

### Mice infection

All animal studies were conducted in compliance with protocols approved by the Northwestern University Institutional Animal Care and Use Committee as previously detailed(*15*). Briefly, ICR mice (Jackson Laboratories, *n*=10 per group) were anaesthetized with a ketamine-xylazine cocktail and then 50 µl 0.9% sodium bicarbonate immediately followed by a 50 µl bacterial suspension (3.5 - 4 x10^7^ CFU/mL PBS) was injected to the stomach by oral gavage using a 1-cm animal feeding needle. Mice were monitored every 6 hours for a total experimental duration of 36 hours post-infection. Surviving mice were euthanized after 48 hours and counted as survivors.

### Cytotoxicity assay

Cells were seeded in a 12-well plate overnight. Before bacteria co-incubation, cells were washed and the medium was replaced with antibiotic-free, serum-free, phenol red-free DMEM media (Gibco, Cat.#31053-028). Cells were then co-incubated with the specified bacterial strains grown to mid-log phase at the indicated multiplicity of infection (MOI) for 180 minutes. Following co-incubation, 20 µg/ml Gentamicin was added, and the supernatants from the spent media were collecte and centrifugated at 13,000 xg for 1 minute. LDH activity was measured by CytoTox 96® Non-Radioactive Cytotoxicity Assay kit (Promega, Cat #G1780) according to the manufacturer’s instructions.

### Generation of *V. vulnificus* strains

Genetic fragments corresponding to the sequences upstream and downstream of the Aknot locus were synthesized as Gblock by Integrated DNA Technologies (Coralville, IA). These fragments were then cloned into the pKEK2200 vector through the Gibson Assembly method (New England Biolabs, Cat.#E2611) following the manufacturer’s protocol. The recombinant plasmid was subsequently introduced into *E.coli* strain SM10λpir and transferred to *V. vulnificus* CMCP6Δ*vvhA* by conjugation, followed by selection of double homologous recombination events through sucrose counterselection as previously described(*59*).

### Affinity purification mass spectrometry

HeLa cells cultured to 80% confluency in a 6-well plate were treated with 500 nM of recombinant Aknot-HA in Hank’s Balanced Salt Solution (HBSS) at 4°C for one hour. Following incubation, cell were lysed using RIPA buffer complemented with protease inhibitor cocktails. The lysates was centrifuged at 13,000 x *g* for 15 minutes and the clarified supernatant was applied to a His SpinTrap column (Cytiva, Cat.#28401353), which had been previously equilibrated with a binding buffer (10 mM Tris-HCl, pH 8.3, 500 mM NaCl). The clarified cell lysate was incubated with Ni^2+^ affinity resins within the column at 4°C overnight. To reduce nonspecific binding, the column was washed with 10 column volumes (CV) of binding buffer and followed by 10 CV of washing buffer containing 10 mM Tris-HCl (pH 8.3), 1 M NaCl, and 25 mM imidazole. Bound proteins were eluted using an elution buffer comprised of 10 mM Tris-HCl (pH 8.3), 500 mM NaCl, and 500 mM imidazole. The eluted fractions were then submitted to Northwestern University Proteomics Center of Excellence Core Facility (NUPCE) for mass spectrometry analysis for protein identifications.

### Establish L1CAM^-/-^ cell line

HeLa L1CAM^-/-^ cells were generated using CRISPR/Cas9 genome editing system. A guide RNA (gRNA) sequence targeting the third exon of the human L1CAM gene (5’-GAGTAGCCGATAGTGACCTG-3’) was designed and cloned into the pKLV2-U6gRNA(BbsI)-PGKpuro2ABFP vector (Addgene plasmid #67974). For the production of lentiviral particles carrying the CRISPR/Cas9 components, HEK293T cells were co-transfected with the constructed pKLV2 gRNA plasmid, the packaging plasmid pAX2, and the envelope plasmid pMD2.G at a ratio of 2:2:1. Transfection was performed using Lipofectamine 2000 (Invitrogen) according to the manufacturer’s protocol. Following transfection, the cells were incubated for 72 hours to allow for viral particle production. The supernatants containing the lentiviral particles were then harvested and filtered through a 0.45 µm filter. HeLa Cas9 cells were infected with the lentiviral particles carrying the CRISPR/Cas9 components. Following 24 hours of infection, the medium was replaced with fresh growth medium containing 2 µg/ml puromycin to select for successfully transduced cells. The selection process was continued for 10 days, cells were diluted and plated at a density to allow for the isolation of individual clones. Single cell-derived colonies were manually picked for isolation and protein levels of L1CAM were determined by immunoblotting to confirm the knockout of L1CAM.

### Statistical analysis

Statistical and graphical analyses were performed using GraphPad Prism version 10 (GraphPad, San Diego, CA, USA). Data were analyzed employing the student’s t-test for comparisons between two groups. The significance levels were denoted as follows: \**p*<0.05, \*\**p*<0.001, and \*\*\**p*<0.0001. Data are presented as mean ± standard deviation (SD).

## Acknowledgements.

We thank Jason A. Pattie for help with AlphaFold2 modeling. This work was funded by NIAID grants R21 AI149061 (to X.W. and K.S.) and R37 AI092825 (to K.S.) and grants from the Danish National Research Foundation (DNRF107), Novo Nordisk Foundation, and Lundbeck Foundation (to Y.N. and H.C.), J.C. was supported by fellowships from the Northwestern Molecular Biophysics Training program (NIGMS T32 GM008382) and NIAID (F31 AI172382). F.G. was supported by a postdoctoral fellowship from the European Molecular Biology Organization (ALTF 105-2023). T.J. was supported by the NEYE Foundation (ID 24030064) and the European Molecular Biology Organization (EMBO) postdoctoral fellowship (ALTF 336-2021), The research used resources of the Northwestern University Interdepartmental Immuno-Biology Flow Cytometry Core Facility, the Center for Advanced Microscopy, the NU Proteomics Center of Excellence Core Facility (NUPCE), and the Structural Biology Facility, which are all supported core facilities of the Robert H. Lurie Comprehensive Cancer Center funded by grant P30CA060553. Additionally, we appreciate the support from the University of Copenhagen Center for Glycomics groups, and the Core Facility for Flow Cytometry and Single Cell Analysis at Faculty of Health and Medical Sciences, University of Copenhagen.

## Author contributions

Conceptualization-JC, XW, HC, KS; Formal Analysis-JC, YN; Investigation-JC, YN, FG, TJ, FXSH; Writing original draft-JC; Writing-review and editing-YN, XW, HC, KS; Visualization-JC; Funding acquisition-JC, HC, KS.

## Data and material availability

All data relevant to this study are found in the manuscript or the supplement. All unique/stable reagents generated in this study are available from Karla Satchell or Henrik Clausen.

## REFERENCES

1. J.-M. Cavaillon, Exotoxins and endotoxins: Inducers of inflammatory cytokines. Toxicon 149, 45–53 (2018).

2. J. K. Actor, “11 - Basic Bacteriology” in Elsevier’s Integrated Review Immunology and Microbiology (Second Edition), J. K. Actor, Ed. (W.B. Saunders, Philadelphia, 2012; https://www.sciencedirect.com/science/article/pii/B9780323074476000119), pp. 93–103.

3. C. K. Schmitt, K. C. Meysick, A. D. O’Brien, Bacterial Toxins: Friends or Foes? - Volume 5, Number 2—April 1999 - Emerging Infectious Diseases journal - CDC. doi: 10.3201/eid0502.990206.

4. C. Ghazaei, Advances in the Study of Bacterial Toxins, Their Roles and Mechanisms in Pathogenesis. Malays J Med Sci 29, 4–17 (2022).

5. B. S. Kim, The Modes of Action of MARTX Toxin Effector Domains. Toxins (Basel) 10, 507 (2018).

6. K. J. F. Satchell, MARTX, multifunctional autoprocessing repeats-in-toxin toxins. Infect Immun 75, 5079–5084 (2007).

7. H. E. Gavin, K. J. F. Satchell, MARTX toxins as effector delivery platforms. Pathog Dis 73, ftv092 (2015).

8. P. J. Woida, K. J. F. Satchell, Coordinated delivery and function of bacterial MARTX toxin effectors. Mol Microbiol 107, 133–141 (2018).

9. K. J. F. Satchell, Structure and function of MARTX toxins and other large repetitive RTX proteins. Annu Rev Microbiol 65, 71–90 (2011).

10. V. Olivier, G. K. Haines, Y. Tan, K. J. F. Satchell, Hemolysin and the multifunctional autoprocessing RTX toxin are virulence factors during intestinal infection of mice with Vibrio cholerae El Tor O1 strains. Infect Immun 75, 5035–5042 (2007).

11. V. Olivier, N. H. Salzman, K. J. F. Satchell, Prolonged colonization of mice by Vibrio cholerae El Tor O1 depends on accessory toxins. Infect Immun 75, 5043–5051 (2007).

12. P. J. Woida, K. J. F. Satchell, The Vibrio cholerae MARTX toxin silences the inflammatory response to cytoskeletal damage before inducing actin cytoskeleton collapse. Sci Signal 13, eaaw9447 (2020).

13. J. S. Kwak, H.-G. Jeong, K. J. F. Satchell, Vibrio vulnificus rtxA1 gene recombination generates toxin variants with altered potency during intestinal infection. Proc Natl Acad Sci U S A 108, 1645–1650 (2011).

14. H.-G. Jeong, K. J. F. Satchell, Additive function of Vibrio vulnificus MARTX(Vv) and VvhA cytolysins promotes rapid growth and epithelial tissue necrosis during intestinal infection. PLoS Pathog 8, e1002581 (2012).

15. H. E. Gavin, N. T. Beubier, K. J. F. Satchell, The Effector Domain Region of the Vibrio vulnificus MARTX Toxin Confers Biphasic Epithelial Barrier Disruption and Is Essential for Systemic Spread from the Intestine. PLoS Pathog 13, e1006119 (2017).

16. G. Suarez, B. K. Khajanchi, J. C. Sierra, T. E. Erova, J. Sha, A. K. Chopra, Actin cross-linking domain of *Aeromonas hydrophila* repeat in toxin A (RtxA) induces host cell rounding and apoptosis. Gene 506, 369–376 (2012).

17. K. J. F. Satchell, Multifunctional-autoprocessing repeats-in-toxin (MARTX) Toxins of Vibrios. Microbiol Spectr 3, VE-001-2014 (2015).

18. C. Baker-Austin, J. D. Oliver, Vibrio vulnificus. Trends in Microbiology 28, 81–82 (2020).

19. M. A. Horseman, S. Surani, A comprehensive review of Vibrio vulnificus: an important cause of severe sepsis and skin and soft-tissue infection. Int J Infect Dis 15, e157–166 (2011).

20. S.-R. Chiang, Y.-C. Chuang, Vibrio vulnificus infection: clinical manifestations, pathogenesis, and antimicrobial therapy. J Microbiol Immunol Infect 36, 81–88 (2003).

21. Health Alert Network (HAN) - 00497 | Severe Vibrio vulnificus Infections in the United States Associated with Warming Coastal Waters (2023). https://emergency.cdc.gov/han/2023/han00497.asp.

22. C. Baker-Austin, J. A. Trinanes, N. G. H. Taylor, R. Hartnell, A. Siitonen, J. Martinez-Urtaza, Emerging Vibrio risk at high latitudes in response to ocean warming. Nature Clim Change 3, 73–77 (2013).

23. E. J. Archer, C. Baker-Austin, T. J. Osborn, N. R. Jones, J. Martínez-Urtaza, J. Trinanes, J. D. Oliver, F. J. C. González, I. R. Lake, Climate warming and increasing Vibrio vulnificus infections in North America. Sci Rep 13, 3893 (2023).

24. R. Deeb, D. Tufford, G. I. Scott, J. G. Moore, K. Dow, Impact of Climate Change on Vibrio vulnificus Abundance and Exposure Risk. Estuaries Coast 41, 2289–2303 (2018).

25. H.-R. Lo, J.-H. Lin, Y.-H. Chen, C.-L. Chen, C.-P. Shao, Y.-C. Lai, L.-I. Hor, RTX Toxin Enhances the Survival of Vibrio vulnificus During Infection by Protecting the Organism From Phagocytosis. The Journal of Infectious Diseases 203, 1866–1874 (2011).

26. K. J. Ziolo, H.-G. Jeong, J. S. Kwak, S. Yang, R. M. Lavker, K. J. F. Satchell, Vibrio vulnificus biotype 3 multifunctional autoprocessing RTX toxin is an adenylate cyclase toxin essential for virulence in mice. Infect Immun 82, 2148–2157 (2014).

27. B. K. Boardman, K. J. F. Satchell, Vibrio cholerae strains with mutations in an atypical type I secretion system accumulate RTX toxin intracellularly. J. Bacteriol. 186, 8137– 8143 (2004).

28. M. Egerer, K. J. F. Satchell, Inositol hexakisphosphate-induced autoprocessing of large bacterial protein toxins. PLoS Pathog 6, e1000942 (2010).

29. D. S. Kudryashov, C. L. Cordero, E. Reisler, K. J. F. Satchell, Characterization of the Enzymatic Activity of the Actin Cross-linking Domain from the *Vibrio cholerae* MARTX*Vc* Toxin*. J Biol Chem 283, 445–452 (2008).

30. P. J. Woida, K. J. F. Satchell, The Vibrio cholerae MARTX toxin silences the inflammatory response to cytoskeletal damage before inducing actin cytoskeleton collapse | Science Signal 13, eaaw9447 (2020)

31. Y. Xu, K. Ding, T. Peng, Chemical Proteomics Reveals Nε-Fatty-Acylation of Septins by Rho Inactivation Domain (RID) of the Vibrio MARTX Toxin to Alter Septin Localization and Organization. Mol Cell Proteomics 23, 100730 (2024).

32. C. L. Cordero, D. S. Kudryashov, E. Reisler, K. J. F. Satchell, The Actin cross-linking domain of the Vibrio cholerae RTX toxin directly catalyzes the covalent cross-linking of actin. J Biol Chem 281, 32366–32374 (2006).

33. K.-L. Sheahan, C. L. Cordero, K. J. F. Satchell, Identification of a domain within the multifunctional Vibrio cholerae RTX toxin that covalently cross-links actin. Proc Natl Acad Sci U S A 101, 9798–9803 (2004).

34. S. Agarwal, Y. Zhu, D. R. Gius, K. J. F. Satchell, The Makes Caterpillars Floppy (MCF)-Like Domain of Vibrio vulnificus Induces Mitochondrion-Mediated Apoptosis. Infect Immun 83, 4392–4403 (2015).

35. A. Herrera, M. M. Packer, M. Rosas-Lemus, G. Minasov, J. Chen, J. H. Brumell, K. J. F. Satchell, Vibrio MARTX toxin processing and degradation of cellular Rab GTPases by the cytotoxic effector Makes Caterpillars Floppy. Proc Natl Acad Sci USA 121, e2316143121 (2024).

36. M. Biancucci, G. Minasov, A. Banerjee, A. Herrera, P. J. Woida, M. B. Kieffer, L. Bindu, M. Abreu-Blanco, W. F. Anderson, V. Gaponenko, A. G. Stephen, M. Holderfield, K. J. F. Satchell, The bacterial Ras/Rap1 site-specific endopeptidase RRSP cleaves Ras through an atypical mechanism to disrupt Ras-ERK signaling. Sci Signal 11, eaat8335 (2018).

37. B. A. Kim, J. Y. Lim, J. H. Rhee, Y. R. Kim, Characterization of Prohibitin 1 as a Host Partner of Vibrio vulnificus RtxA1 Toxin. J Infect Dis 213, 131–138 (2016).

38. Y.-T. Peng, P. Chen, R.-Y. Ouyang, L. Song, Multifaceted role of prohibitin in cell survival and apoptosis. Apoptosis 20, 1135–1149 (2015).

39. B. S. Kim, H. E. Gavin, K. J. F. Satchell, Distinct roles of the repeat-containing regions and effector domains of the Vibrio vulnificus multifunctional-autoprocessing repeats-in-toxin (MARTX) toxin. mBio 6 (2015).

40. J. Jumper, R. Evans, A. Pritzel, T. Green, M. Figurnov, O. Ronneberger, K. Tunyasuvunakool, R. Bates, A. Žídek, A. Potapenko, A. Bridgland, C. Meyer, S. A. A. Kohl, A. J. Ballard, A. Cowie, B. Romera-Paredes, S. Nikolov, R. Jain, J. Adler, T. Back, S. Petersen, D. Reiman, E. Clancy, M. Zielinski, M. Steinegger, M. Pacholska, T. Berghammer, S. Bodenstein, D. Silver, O. Vinyals, A. W. Senior, K. Kavukcuoglu, P. Kohli, D. Hassabis, Highly accurate protein structure prediction with AlphaFold. Nature 596, 583–589 (2021).

41. A. N. Barclay, Membrane proteins with immunoglobulin-like domains—a master superfamily of interaction molecules. Seminars in Immunology 15, 215–223 (2003).

42. N. E. Sanjana, O. Shalem, F. Zhang, Improved vectors and genome-wide libraries for CRISPR screening. Nat Methods 11, 783–784 (2014).

43. D. Medina-Cano, E. Ucuncu, L. S. Nguyen, M. Nicouleau, J. Lipecka, J.-C. Bizot, C. Thiel, F. Foulquier, N. Lefort, C. Faivre-Sarrailh, L. Colleaux, I. C. Guerrera, V. Cantagrel, High N-glycan multiplicity is critical for neuronal adhesion and sensitizes the developing cerebellum to N-glycosylation defect. eLife 7, e38309 (2018).

44. Y. Narimatsu, H. J. Joshi, R. Nason, J. Van Coillie, R. Karlsson, L. Sun, Z. Ye, Y.-H. Chen, K. T. Schjoldager, C. Steentoft, S. Furukawa, B. A. Bensing, P. M. Sullam, A. J. Thompson, J. C. Paulson, C. Büll, G. J. Adema, U. Mandel, L. Hansen, E. P. Bennett, A. Varki, S. Y. Vakhrushev, Z. Yang, H. Clausen, An Atlas of Human Glycosylation Pathways Enables Display of the Human Glycome by Gene Engineered Cells. Molecular Cell 75, 394–407.e5 (2019).

45. C. Büll, H. J. Joshi, H. Clausen, Y. Narimatsu, Cell-Based Glycan Arrays-A Practical Guide to Dissect the Human Glycome. STAR Protoc 1, 100017 (2020).

46. H. Miller-Podraza, K. Weikkolainen, T. Larsson, P. Linde, J. Helin, J. Natunen, K.-A. Karlsson, Helicobacter pylori binding to new glycans based on N-acetyllactosamine. Glycobiology 19, 399–407 (2009).

47. M. P. Jennings, C. J. Day, J. M. Atack, How bacteria utilize sialic acid during interactions with the host: snip, snatch, dispatch, match and attach. Microbiology 168, 001157 (2022).

48. T, Jaroentomeechai, et al, Mammalian Cell-Based Production of Glycans, Glycopeptides and Glycomodules - Expanding the Glycoscience Toolbox. Nat Method (In review).

49. M. Uhlen, et al, Proteomics. Tissue-based map of the human proteome. Science 347, 394 (2015).

50. K. Ganesh, H. Basnet, Y. Kaygusuz, A. M. Laughney, L. He, R. Sharma, K. P. O’Rourke, V. P. Reuter, Y.-H. Huang, M. Turkekul, E. E. Er, I. Masilionis, K. Manova-Todorova, M. R. Weiser, L. B. Saltz, J. Garcia-Aguilar, R. Koche, S. W. Lowe, D. Pe’er, J. Shia, J. Massagué, L1CAM defines the regenerative origin of metastasis-initiating cells in colorectal cancer. Nat Cancer 1, 28–45 (2020).

51. N. Song, L. Chen, X. Ren, N. R. Waterfield, J. Yang, G. Yang, N-Glycans and sulfated glycosaminoglycans contribute to the action of diverse Tc toxins on mammalian cells. PLOS Pathogens 17, e1009244 (2021).

52. P. N. Ng’ang’a, L. Siukstaite, A. E. Lang, H. Bakker, W. Römer, K. Aktories, G. Schmidt, Involvement of N-glycans in binding of Photorhabdus luminescens Tc toxin. Cellular Microbiology 23, e13326 (2021).

53. D. Roderer, F. Bröcker, O. Sitsel, P. Kaplonek, F. Leidreiter, P. H. Seeberger, S. Raunser, Glycan-dependent cell adhesion mechanism of Tc toxins. Nat Commun 11, 2694 (2020).

54. S. J. Piper, L. Brillault, R. Rothnagel, T. I. Croll, J. K. Box, I. Chassagnon, S. Scherer, K. N. Goldie, S. A. Jones, F. Schepers, L. Hartley-Tassell, T. Ve, J. N. Busby, J. E. Dalziel, J. S. Lott, B. Hankamer, H. Stahlberg, M. R. H. Hurst, M. J. Landsberg, Cryo-EM structures of the pore-forming A subunit from the Yersinia entomophaga ABC toxin. Nat Commun 10, 1952 (2019).

55. E. N. Wallace, C. A. West, C. T. McDowell, X. Lu, E. Bruner, A. S. Mehta, K. F. Aoki-Kinoshita, P. M. Angel, R. R. Drake, An N-glycome tissue atlas of 15 human normal and cancer tissue types determined by MALDI-imaging mass spectrometry. Sci Rep 14, 489 (2024).

56. G. E. Seabright, C. A. Cottrell, M. J. van Gils, A. D’addabbo, D. J. Harvey, A.-J. Behrens, J. D. Allen, Y. Watanabe, N. Scaringi, T. M. Polveroni, A. Maker, S. Vasiljevic, N. de Val, R. W. Sanders, A. B. Ward, M. Crispin, Networks of HIV-1 Envelope Glycans Maintain Antibody Epitopes in the Face of Glycan Additions and Deletions. Structure 28, 897–909.e6 (2020).

57. J. Luo, Q. Yang, X. Zhang, Y. Zhang, L. Wan, X. Zhan, Y. Zhou, L. He, D. Li, D. Jin, Y. Zhen, J. Huang, Y. Li, L. Tao, TFPI is a colonic crypt receptor for TcdB from hypervirulent clade 2 C. difficile. Cell 185, 980–994.e15 (2022).

58. Z. Yang, S. Wang, A. Halim, M. A. Schulz, M. Frodin, S. H. Rahman, M. B. Vester-Christensen, C. Behrens, C. Kristensen, S. Y. Vakhrushev, E. P. Bennett, H. H. Wandall, H. Clausen, Engineered CHO cells for production of diverse, homogeneous glycoproteins. Nat Biotechnol 33, 842–844 (2015).

59. K. J. Fullner, J. J. Mekalanos Genetic characterization of a new type IV-A pilus gene cluster found in both classical and El Tor biotypes of Vibrio cholerae. Infection and immunity 67, 1393–1404 (1999).

