## Supplemental Figures for "Biantennary N-glycans As Receptors for MARTX Toxins in *Vibrio* Pathogenesis"

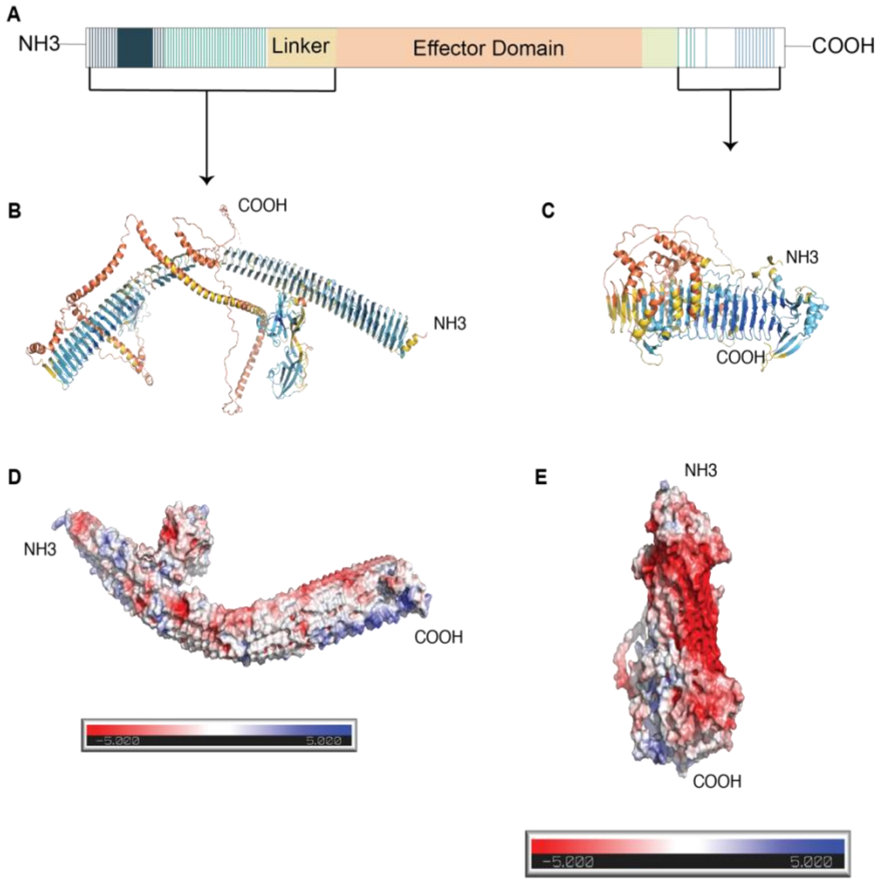


**Fig. S1 AlphaFold structural predictions of MARTX$_{Vv}$:** (A) Schematic representation of the structure organization of the *V. vulnificus* MARTX toxin. (B) An AlphaFold2 structural prediction of the N-terminal regions of MARTX$_{Vv,}$ spanning from "A" repeat to the linker region. (C) An AlphaFold2 structural prediction of the C-terminal repeat regions of MARTX$_{Vv.}$ The structures are color-coded based on the pLDDT scores. (D) Electrostatic projection of the first 1389 aa of MARTX$_{Vv}$ ("A" + "B" repeats). (E) Electrostatic projection of the C-terminal repeat regions of MARTX$_{Vv}$.

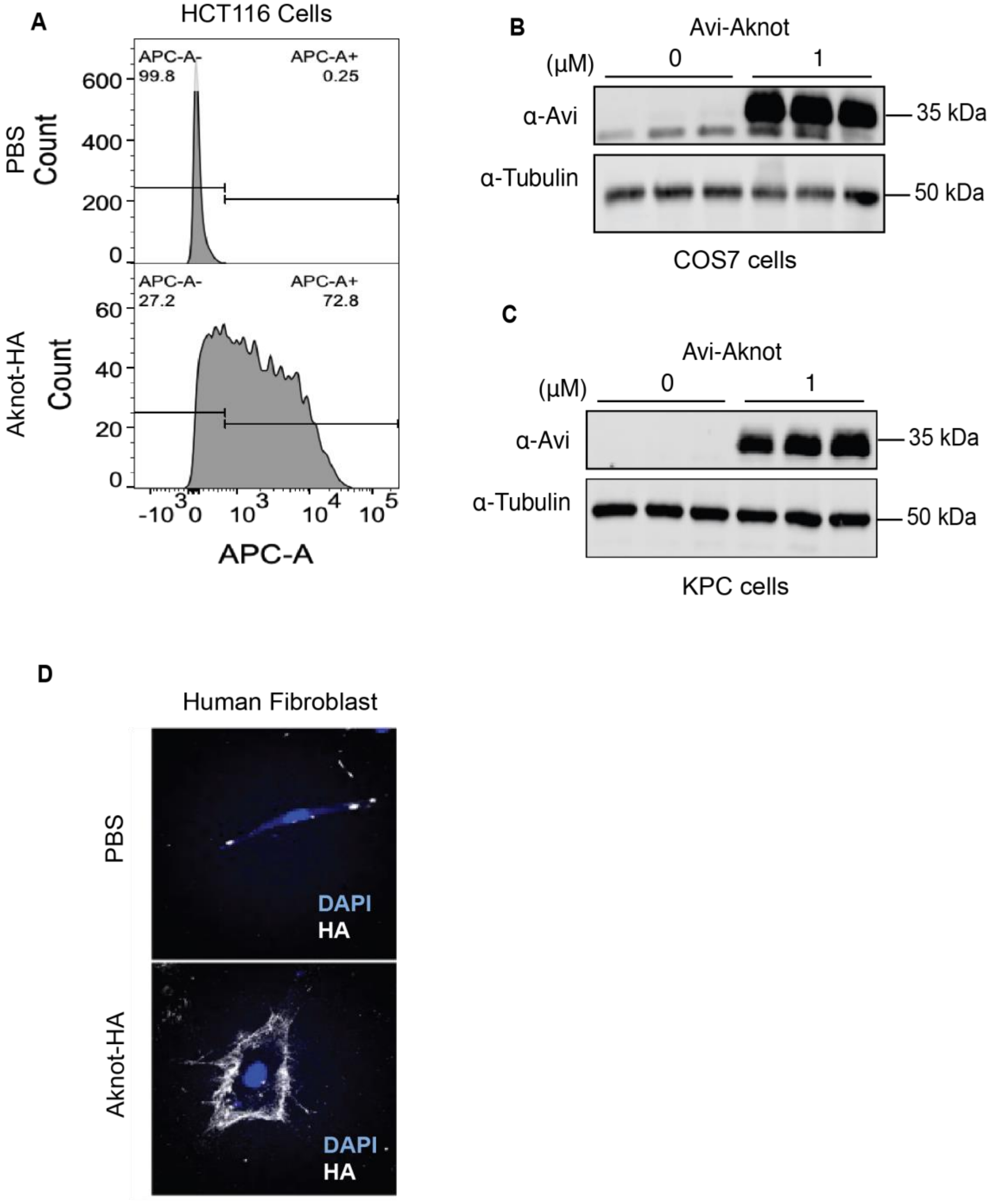


**Fig. S2 Aknot binds to multiple cell lines.** (A) Flow cytometry analysis of 100 nM of Aknot-HA binding to intestinal HCT116 cells. (B) Representative western blot of N-terminal Avi-tagged Aknot (Avi-Aknot) binding to African Green monkey kidney fibroblast-like COS7 cells at indicated concentrations. (C) Western blot analysis of (Avi-Aknot) binding to mouse pancreatic KPC cells at indicated concentrations. (D) Immunofluorescence imaging of Aknot-HA binding to human fibroblast cells.

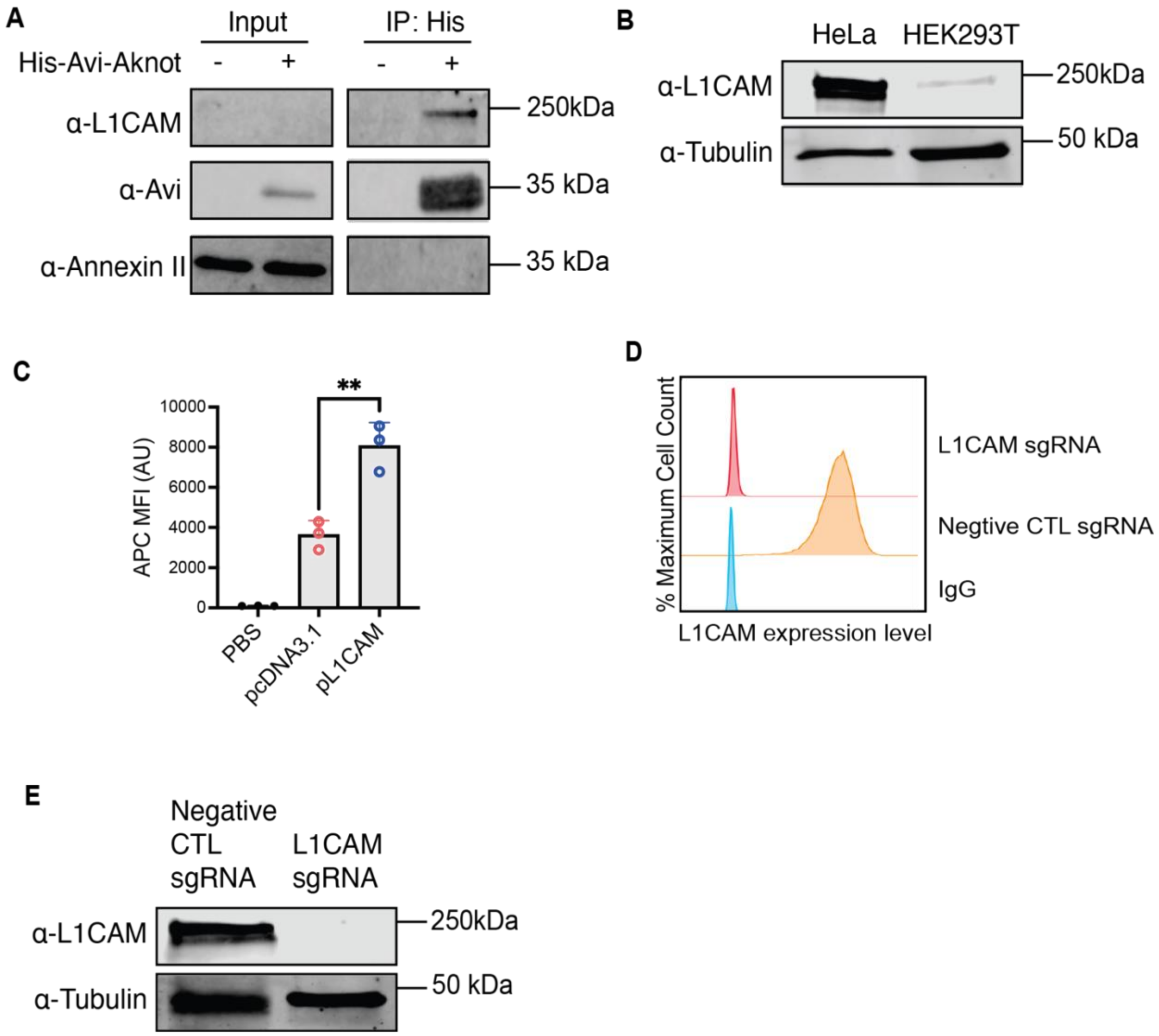


**Fig. S3 Aknot interacts with cell surface L1CAM.** (A) Co-immunoprecipitation analysis of Aknot-HA with L1CAM. HeLa cells were treated with 500 nM 6xHis-Avi-Aknot and subjected to co-immunoprecipitation with $Ni^{+}$ resin loaded HisSpin columns. Western blotting was conducted to detect L1CAM, Avi-tagged Aknot, as well as Annexin V which serves as a negative control. Due to the limitations imposed by the quantity of cell lysate loaded as input, L1CAM was not detectable. However, its presence in the input fraction was confirmed through its detection in the eluted fractions. (B) Western blot analysis of L1CAM expression in HeLa and HEK293T cells. (C) Quantifications of flow cytometry analysis of Aknot-HA binding to HEK293T cells transfected with either an empty vector (pcDNA3.1) or the vector encoding the L1CAM sequence (pL1CAM). Data were represented as mean ± SD from *n*=3 individual experiments (**p<0.001, Student's *t* test). (D) Flow cytometry analysis of the expression level of L1CAM on cell surface on HeLa Cas9 $L1CAM^{-/-}$ and its control cells. (E) Western blot analysis of L1CAM expression in HeLa Cas9 $L1CAM^{-/-}$ and its control cells.

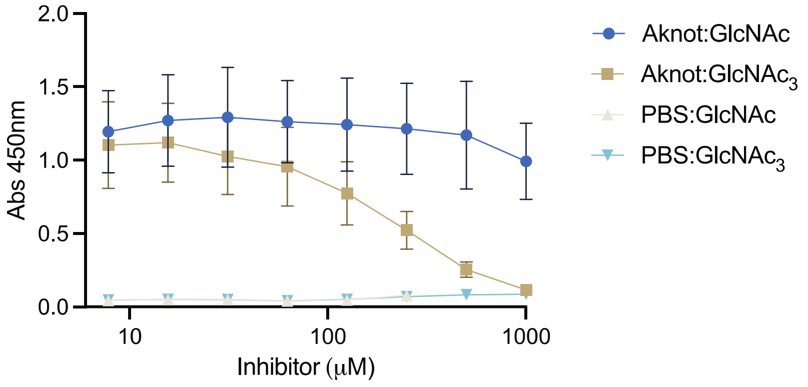


**Fig. S4 Aknot interacts with GlcNAc:** ELISA binding curve of Aknot-HA to the ectodomain of L1CAM. Aknot-HA was pre-incubated with GlcNAc or $GlcNAc_3$ at indicated concentrations, and then was subjected to binding to ELISA plates coated with L1CAM.

**Table S1.**

| Aknot interacting with L1CAM identified by affinity purification mass spectrometry | | | | |
|---|---|---|---|---|
| Protein | Molecular Weight | Percent<br><br>Coverage | # of unique peptides | Exp# |
| L1CAM | 140 kDa | 3.60% | 4 | 1 |
| L1CAM | 140 kDa | 9.60% | 11 | 2 |

**Data S1. (separate file)**

Affinity purification mass spectrometry (AP-MS) identified peptides

**Data S2. (separate file)**

Consortium for Functional Glycomics (CFG) mammalian-type glycan microarrays screening
